# Suppression of YAP Safeguards Human Naïve Pluripotency

**DOI:** 10.1101/2022.06.13.495936

**Authors:** Anish Dattani, Tao Huang, Austin Smith, Ge Guo

**Affiliations:** Living Systems Institute, University of Exeter, Stocker Road, Exeter EX4 4QD, United Kingdom

## Abstract

Propagation of human naïve pluripotent stem cells (nPSCs) requires inhibition of MEK/ERK signalling. However, MEK/ERK inhibition also induces differentiation into trophectoderm (TE). Therefore, robust self-renewal requires active suppression of TE fate. Tankyrase inhibition using XAV939 has been shown to stabilise human nPSCs. Here we dissect the mechanism of this effect. Tankyrase inhibition blocks canonical Wnt/β-catenin signalling. However, nPSCs depleted of β-catenin remain dependent on XAV939. We show that XAV939 prevents TE induction by suppressing YAP activity independent of β-catenin. Tankyrase inhibition stabilises angiomotin, which reduces nuclear translocation of YAP1/TAZ. Upon deletion of Angiomotin-family members *AMOT* and *AMOTL2*, nuclear YAP increases and XAV939 fails to prevent TE induction. Conversely, nPSCs lacking YAP1 fail to undergo TE differentiation and sustain efficient self-renewal without XAV939. These findings explain the distinct requirement for tankyrase inhibition in human but not mouse naïve PSCs and highlight the pivotal role of YAP in human naïve pluripotency.

## INTRODUCTION

Pluripotent stem cells (PSC) are a unique resource for human developmental biology as well as a powerful system for biomedical research. Pluripotent stem cells related to the naïve epiblast in the pre-implantation embryo were first established from mice (Evans and Kaufman, 1981; Martin, 1981). Mouse embryonic stem (ES) cells can be propagated in a highly homogeneous condition in defined media comprising inhibitors of the mitogen-activated protein kinase (ERK1/2) pathway and of glycogen synthase kinase 3 (GSK3) together with the cytokine Leukaemia inhibitory factor (LIF), a formula termed 2iLIF (Ying et al., 2008). However, 2iLIF proved insufficient to capture human naïve PSCs (nPSCs). Two derivative culture conditions were later identified that, in combination with mouse feeder cells, support human nPSCs exhibiting transcriptome proximity to pre-implantation epiblast (Takashima et al., 2014; Theunissen et al., 2014). These formulae (5iLA and t2iLGö) contained the MEK/ERK inhibitor, PD0325901 and the GSK3 inhibitor CHIR99021 with additional kinase inhibitors plus LIF. Subsequently it was shown that human nPSCs can be established and robustly expanded without GSK3 inhibition using PD0325901 and LIF together with the aPKC inhibitor Gö6983 and the tankyrase inhibitor XAV939, a condition termed PXGL (Bredenkamp et al., 2019b; Guo et al., 2017).

The difference in self-renewal requirements for mouse and human nPSCs may be related to their differing lineage potency. In both mouse and human nPSCs MEK/ERK inhibition sustains naïve pluripotency by impeding progression to the formative stage of pluripotency (Kalkan et al., 2019; Kunath et al., 2007; Rostovskaya et al., 2019; Smith, 2017). However, unlike mouse ES cells, human naïve PSCs can also differentiate into trophectoderm (TE) (Guo et al., 2021; Io et al., 2021) the first extraembryonic lineage in the mammalian embryo. Remarkably, inhibition of MEK/ERK directly promotes TE induction. This effect must be countermanded to sustain human nPSCs. We previously noted that XAV939 suppresses TE, but the exact mechanism is not clear (Guo et al., 2021).

XAV939 was originally identified as a Wnt pathway inhibitor in a chemical genetic screen (Huang et al., 2009). It is a selective inhibitor of the enzymatic activity of tankyrases 1 and 2 (TNKS1, TNKS2). Tankyrases add poly-ADP-ribose to proteins, leading to degradation by the ubiquitin proteasome pathway (Smith et al., 1998). A prominent tankyrase substrate is AXIN, the scaffold protein in the β-catenin destruction complex. By stabilising AXIN, tankyrase inhibition promotes degradation of β-catenin, the central effector of canonical Wnt signalling (Huang et al., 2009). However, tankyrases have other targets which include the Angiomotin protein family (AMOT) (Bhardwaj et al., 2017; Wang et al., 2018a). AMOT proteins attenuate YAP1/TAZ nuclear translocation by promoting the kinase activity of LATS1/2 (Hirate et al., 2013; Zhao et al., 2011). Tankyrase inhibition can thus impede YAP1/TAZ activity, simulating HIPPO pathway activity (Wang et al., 2015; Zhao et al., 2011). HIPPO and the AMOT/YAP axis play a well characterised role in regulating the segregation of TE and inner cell mass (ICM) in the early mouse embryo (Hirate et al., 2013; Nishioka et al., 2009). Here, we examine the actors downstream of XAV939 in human naïve stem cell maintenance and TE differentiation.

## RESULTS

### Deletion of β-catenin does not alter naïve PSC dependency on tankyrase inhibition

Tankyrase inhibition is commonly employed in media for propagating pluripotent stem cells corresponding to formative or gastrulation stage epiblast in order to prevent primitive streak-like induction by Wnt/β-catenin (Kim et al., 2013; Kinoshita et al., 2021; Kojima et al., 2014; Sumi et al., 2013; Tsakiridis et al., 2014). However, for human nPSCs the contribution of XAV939 to stable expansion in PXGL is primarily related to suppression of TE. Withdrawal of XAV939 from PXGL leads to expression of TE markers *GATA3* and *HAVCR1* (Guo et al., 2021) whereas the Wnt/β-catenin targets *TBXT and MIXL1* are only up-regulated when cells are also released from MEK/ERK inhibition (**Fig. 1A**). Nonetheless we investigated whether genetic ablation of β-catenin suppressed TE induction. We mutated *CTNNB1*, the β-catenin coding gene, in Cas9 expressing HNES1-*GATA3:mKO2* reporter cells (hereafter HNES1-GATA3:mKO2/Cas9). A pool of gRNA transfected cells was established by puromycin selection for 4 days then expanded in PXGL (**Figure S1A**). Flow cytometry confirmed absence of β-catenin in ∼70% cells after 10 passages, indicating no major selective disadvantage (**Fig. 1B**). We picked and expanded individual colonies. By immunoblotting and immunohistochemistry, we identified a clone (*CTNNB1* KO) with undetectable β-catenin protein (**Fig. 1C, Fig. S1B**). Addition of CHIR99021, a Wnt agonist, induced upregulation of Wnt target genes *TBXT* and *MIXL1* in parental cells, but not in *CTNNB1* KO cells, confirming functional inactivation of the Wnt/β-catenin pathway (**Fig. 1D**). *CTNNB1* KO cells formed dome-shaped colonies like parental cells, although we noticed that if passaging was delayed for longer than three days cells in larger colonies cells began to disperse (**Fig. S1C)**. This phenotype may reflect a deficiency in cell-cell adhesion, as observed in *Ctnnb1* KO mouse ESCs (Lyashenko et al., 2011; Wray et al., 2011). Nonetheless, with a 3-day passaging regimen *CTNNB1* KO cells could be stably expanded in PXGL without accumulation of differentiated cells. They maintained naïve marker expression (**Fig. S1D, Fig. S1E**) and displayed the cell surface marker phenotype SUSD2+/CD24-that discriminates naïve from primed hPSCs (Bredenkamp et al., 2019a) (**Fig, 1E, Fig. 1F**).

**Figure 1:**
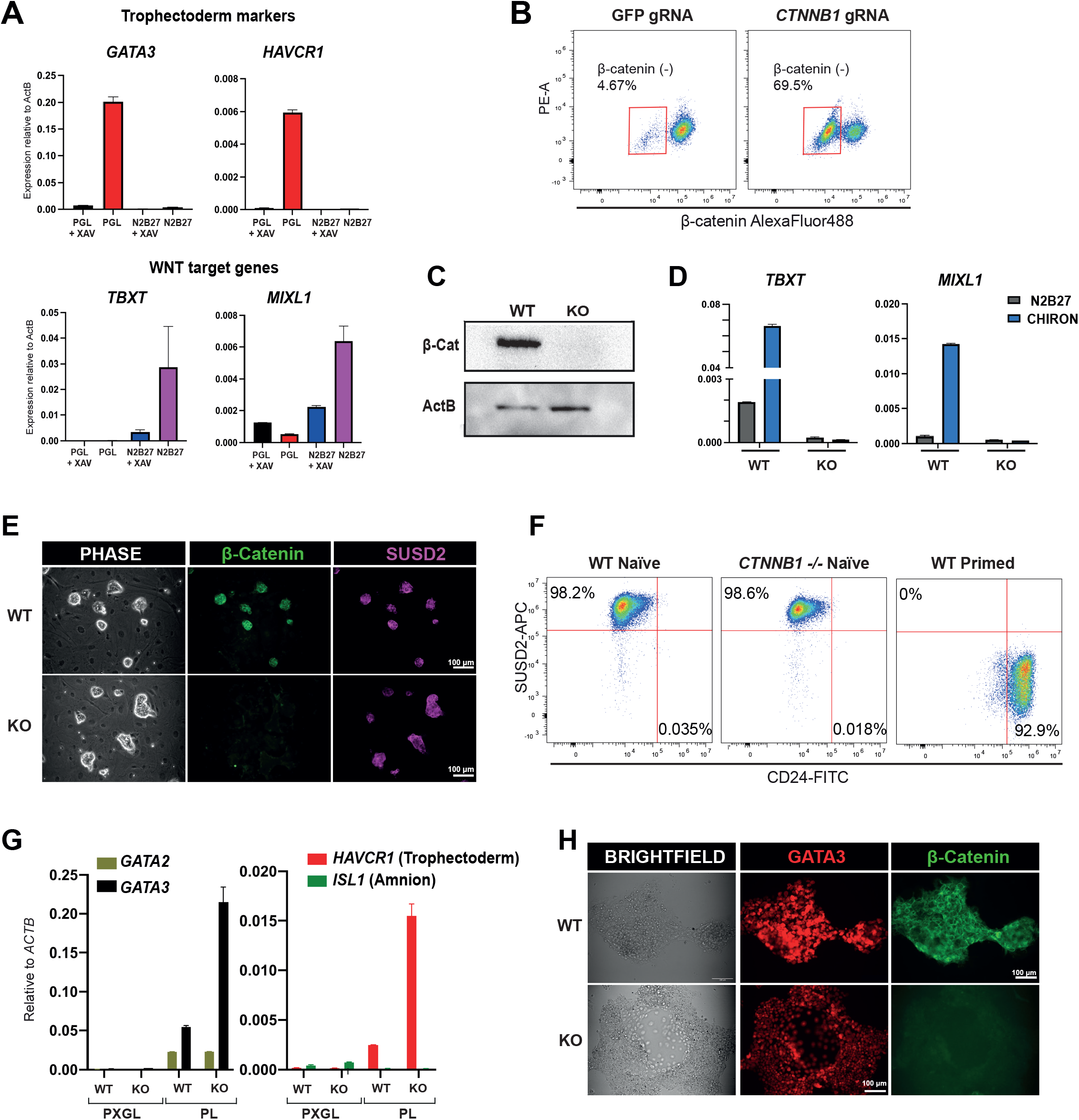
Deletion of β-catenin does not alter naïve PSC dependency on tankyrase inhibition. **A**. qRT-PCR assay for trophectoderm (TE) and Wnt targets genes with or without XAV939 for 3 days, following transfer from PXGL into PGL (PD03, Gö6983, LIF) or N2B27 based medium. Error bars indicate s.d. of PCR duplicates. **B**. Flow cytometry analysis of β-catenin in *CTNNB1* gRNA transfected pool expanded for 10 passages in PXGL. **C**. Western blot for β-catenin in wild-type and clonal *CTNNB1* KO cells. **D**. qRT-PCR for Wnt target genes 24 hours following transfer from PXGL into N2B27 or N2B27 supplemented with 3μM CHIR99021. Error bars indicate s.d. of PCR duplicates. **E**. Phase and immunofluorescence images of wild-type and *CTNNB1* KO cells in PXGL on MEFs. Scale bar: 100 µm. **F**. Flow cytometry analysis of surface markers SUSD2 and CD24 in naïve *HNES1 GATA3:mKO2*/Cas9 (WT Naïve), *CTNNB1* KO and primed HNES1 cells generated by capacitation (WT primed) (Rostovskaya et al., 2019). **G**. qRT-PCR assay for common (*GATA2, GATA3*), TE specific (*HAVCR1*) and Amnion specific (*ISL1*) markers following transfer from PXGL to PL. Error bars indicate s.d. of PCR duplicates. **H**. Phase and immunofluorescence stained images of cells in PL for 3 days. Scale bar: 100 µm.

We investigated TE differentiation in *β*-catenin depleted cells. When the *gRNA* transfectant population or clonal *CTNNB1* KO cells were plated in TE-inductive conditions of PD03 and LIF (PL) without XAV they up-regulated expression of early TE markers *GATA2, GATA3* and *HAVCR1* (**Fig. 1G, Fig. S1F**). Specificity of TE lineage induction was supported by absence of the amnion marker *ISL1* (Zheng et al., 2019). We also observed that *CTNNB1* KO cells readily formed *GATA3:mKO2* positive TE cysts (**Fig. 1H**). We surmise that the formation of epithelial adherens junctions may be sustained by expression of plakoglobin (**Fig. S1G**), which is known to compensate for cell-cell adhesion defects in *Ctnnb1* KO mESCs (Lyashenko et al., 2011).

Overall, these findings establish both that elimination of canonical Wnt signalling does not have a major influence on human nPSC self-renewal, and that β-catenin does not mediate naïve PSC to TE differentiation. We conclude that tankyrase inhibition acts independently of β-catenin destruction to suppress TE fate and sustain human naïve PSC self-renewal.

### XAV939 withdrawal leads to decreased AMOTL2 and upregulation of YAP/TAZ targets

In immortalised cell lines tankyrase inhibition has been shown to stabilise AMOT proteins leading to a reduction in YAP/TAZ nuclear localization and transcriptional co-factor activity (Troilo et al., 2016; Wang et al., 2015). The AMOT family has three members in mammals: AMOT, AMOT-like 1 (AMOTL1) and AMOT-like 2 (AMOTL2). In the early mouse embryo, AMOT and AMOTL2 prevent YAP nuclear localization and TE lineage specification, disabling TEAD-mediated transcription of TE genes in the inner cell mass (ICM) (Hirate et al., 2013; Leung and Zernicka-Goetz, 2013). We surveyed AMOT family expression in published RNA-seq data for human embryos and naïve PSCs (Boroviak et al., 2018; Stirparo et al., 2021). Of the three paralogues, AMOTL2 is most abundant in in nPSCs as well as in the ICM and naïve epiblast of the human embryo (**Fig. 2A, Fig. S2A**), consistent with observations in mouse. We therefore focussed on AMOTL2. The long isoform of AMOTL2 (p100; gene accession no. Q9Y2J4) is associated with cellular junctions. AMOTL2 p100 contains both YAP1 and tankyrase binding domains in the N-terminus and has been shown to be the target of poly-ADP-ribosylation by tankyrase (Cox et al., 2015; Mojallal et al., 2014; Wang et al., 2015) (**Fig. 2B**). We observed reduced levels of AMOTL2 p100 24 hours after XAV removal, whereas expression of the shorter isoform (AMOTL2 p60; gene accession no. AAH11454), which lacks YAP1/TAZ and tankyrase binding domains(Cox et al., 2015) was not significantly changed (**Fig. 2C, Fig. S2C**). Reduced AMOTL2 protein expression was sustained for at least 3 days (**Fig. S2B, Fig. S2D**). We examined YAP1 and TAZ protein localisation by immunofluorescence staining. We noted that YAP1 exhibited nuclear staining with or without XAV (**Fig. 2D**). However, cultures without XAV showed a gradual but significant increase in nuclear YAP1 expression over three days (**Fig. 2E, Fig. 2F**). TAZ was localized in both the nucleus and cytoplasm in PXL cultures (**Fig. 2D**). As for YAP1, withdrawal of XAV lead to increased nuclear TAZ (**Fig. 2D, Fig. 2E**). By 3 days, high nuclear YAP1 and TAZ staining co-localised with GATA3 expression (**Fig. 2F**).

**Figure 2:**
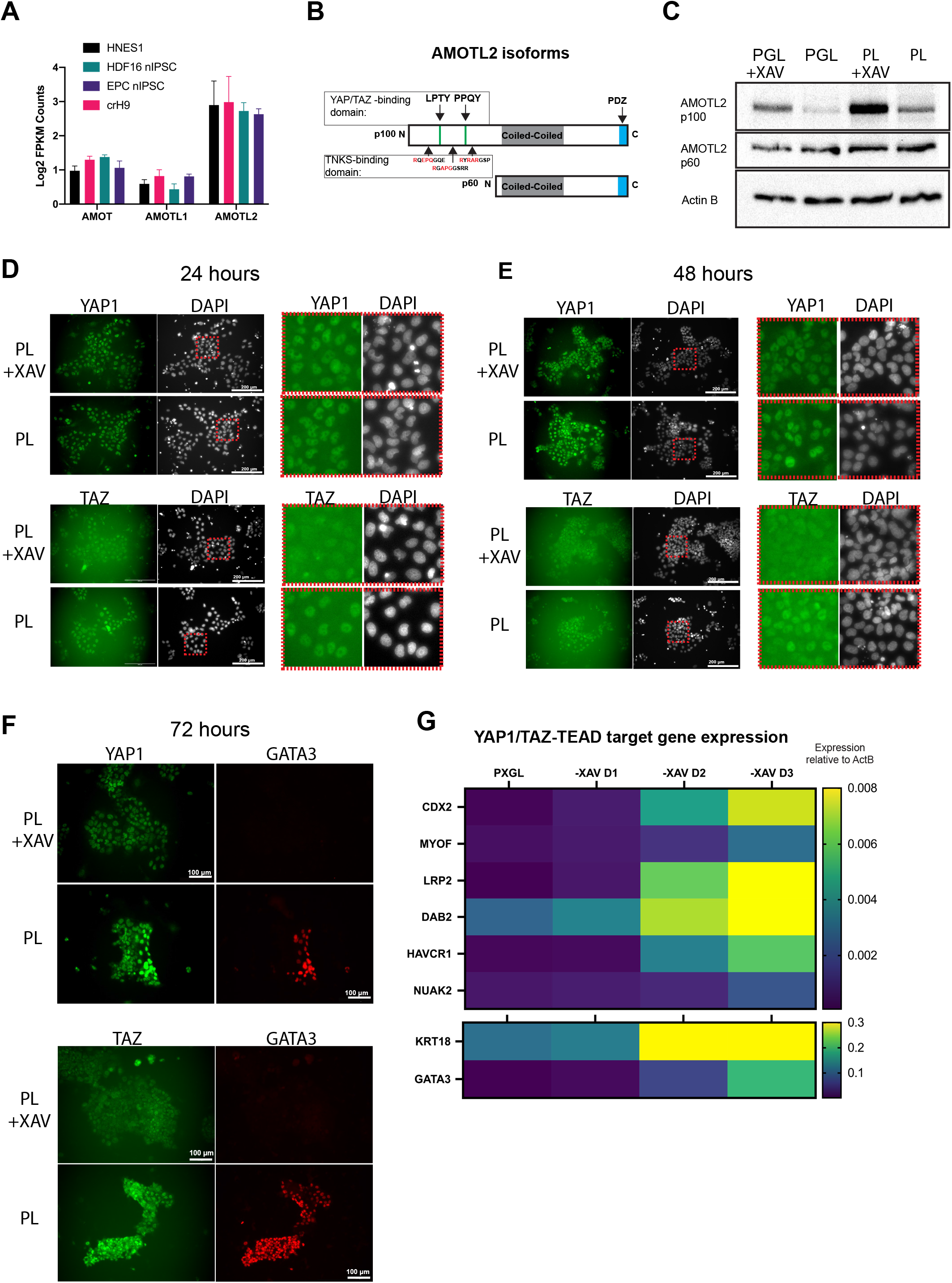
XAV939 withdrawal leads to reduced AMOTL2 protein and upregulation of YAP targets. **A**. FPKM counts for AMOT paralogs in naïve PSCs in published dataset GSE150933 (Stirparo et al., 2021). **B**. Schematic of AMOTL2 long (p100) and short (p60) isoforms with YAP1/TAZ, TNKs and PDZ binding domains **C**. Western blot for AMOTL2 24 hours after transfer from PXGL to indicated conditions (see also Fig. S2B). **D**. Immunofluorescence staining of YAP1 and TAZ (WWTR1) 24 hours after XAV withdrawal from PXL cultures. Scale bar: 200 µm. Red box outlines magnified field in right hand panels. **E**. Immunofluorescence staining of YAP1 and TAZ (WWTR1) 48 hours after XAV withdrawal from PXL cultures. Scale bar: 200 µm. Red box outlines magnified field in right hand panels. **F**. Immunofluorescence staining of YAP1 or TAZ with GATA3 after 72 hrs in PL+XAV or PL. Scale bar: 100 µm. **G**. Heatmap of expression of YAP/TEAD target genes upregulated 24 hrs after transfer from PXGL to PGL. Expression values are means of duplicate qRT-PCR assays.

In the nucleus, YAP1/TAZ acts as a co-factor for TEAD transcription factors (Vassilev et al., 2001). YAP/TEAD target genes have been identified in human cancers(Wang et al., 2018) as well as in mouse trophectoderm development (Posfai et al., 2017). Consistent with increased YAP/TAZ nuclear localization a panel of YAP/TEAD targets are upregulated in nPSCs following XAV removal (**Fig. 2G**). Induction of YAP/TEAD targets in the absence of XAV was also detected in nPSCs generated by somatic cell reprogramming (niPSCs) (**Fig. S2E**).

### XAV inhibition of TE induction is mediated by AMOT proteins

We tested whether the effect of tankyrase inhibition on TE induction requires AMOT. The three *AMOT* genes were mutated individually or in pairwise combination in HNES1-*GATA3: mKO2/Cas9 cells*. We examined *GATA3:mKO2* expression 5 days after gRNA transfection in PXGL. We detected around 10% of cells expressing *GATA3:mKO2* in the *AMOTL2* gRNA transfected pool (**Fig. 3A, Fig. 3B**). Western blotting confirmed reduced AMOTL2 protein in knockout pools (**Fig. S3A)**. Co-transfection of *AMOT* and *AMOTL2* gRNAs yielded a greater increase in *GATA3:mKO2* expression with nearly 40% of cells positive (**Fig. 3A**). AMOTL1 gRNA transfection had a negligible effect (**Fig. S3B**). The expression of *GATA3:mKO2* was accompanied by the up-regulation of TE markers, increased nuclear YAP1 staining, and the expression of YAP/TEAD targets in the presence of XAV939 (**Fig. 3C, Fig. S3C,D**). These results suggest that AMOT family function is required for XAV939-mediated suppression of TE. Of note, however, *AMOT/AMOTL2* depletion only had a marginal effect on GATA3 expression in the absence of PD03 (**Fig. 3D**), suggesting MEK/ERK inhibition is essential for efficient TE induction from human nPSCs.

**Figure 3:**
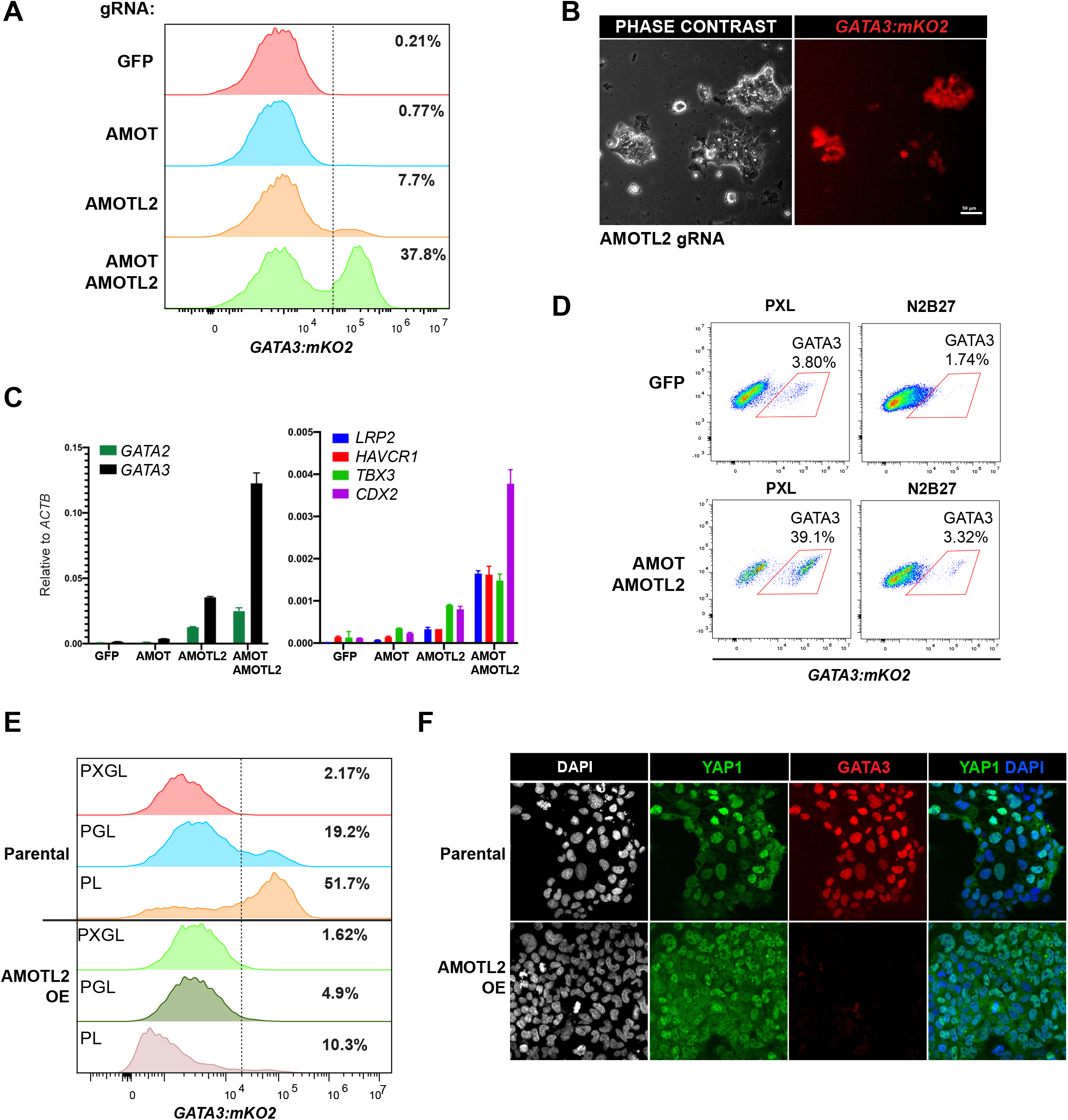
XAV inhibition of TE induction is mediated by AMOT proteins. **A**. Flow cytometry analysis for *GATA3:mKO2* reporter in cells in PXGL 5 days after indicated gRNA transfection and puromycin selection. Histograms show % of maximum events. **B**. Phase and fluorescence imaging showing cells expressing *GATA3:mKO2* reporter in AMOTL2 gRNA transfected cells in PXGL. Scale bar: 50 µm. **C**. qRT-PCR assay for TE markers in cells transfected with indicated gRNAs and cultured in PXGL for 5 days. Error bars indicate s.d. of PCR duplicates. **D**. Flow cytometry for *GATA3:mKO2* reporter expression following AMOT/AMOTL2 double gRNA transfection and culture in PL+XAV or N2B27 for 4 days. **E**. Flow cytometry for *GATA3:mKO2* of parental and AMOTL2 overexpressing (OE) cells transferred to the indicated conditions for 3 days. **F**. Immunofluorescence staining of YAP1 and GATA3 in parental and AMOTL2 OE cells cultured in PL for 3 days.

We then investigated whether overexpression of *AMOTL2* could counteract PD03 induced TE differentiation. Using a CAG expression vector, we constitutively expressed *AMOTL2* in HNES1-*GATA3:mKO2* cells. After removal of XAV we saw little or no expression of *GATA3:mKO2* or TE markers in AMOTL2 overexpressing cells (**Fig. 3E, Fig. S3D**). YAP1 protein was reduced and distributed between the cytoplasm and nucleus in contrast to the nuclear accumulation in parental cells in PD03 (**Fig. 3F**).

These results indicate that AMOT proteins are both necessary and sufficient for the effect of tankyrase inhibition in suppressing TE differentiation.

### Depletion of YAP1 enables sustained self-renewal without XAV939

To confirm the role of YAP/TAZ signalling we deleted *YAP1* or its paralog *TAZ* (*WWTR1*). Pools of cells transfected with either YAP1 or TAZ gRNA showed markedly reduced frequency of GATA3:mKO2 induction when transferred to PD03 (**Fig. 4A**). Co-transfection of *YAP1* and *TAZ* gRNAs completely abolished *GATA3:mKO2* induction. qRT-PCR analysis confirmed reduced TE marker expression in *YAP1, TAZ* and *YAP1/TAZ* co-depleted cells (**Fig. 4B**).

**Figure 4:**
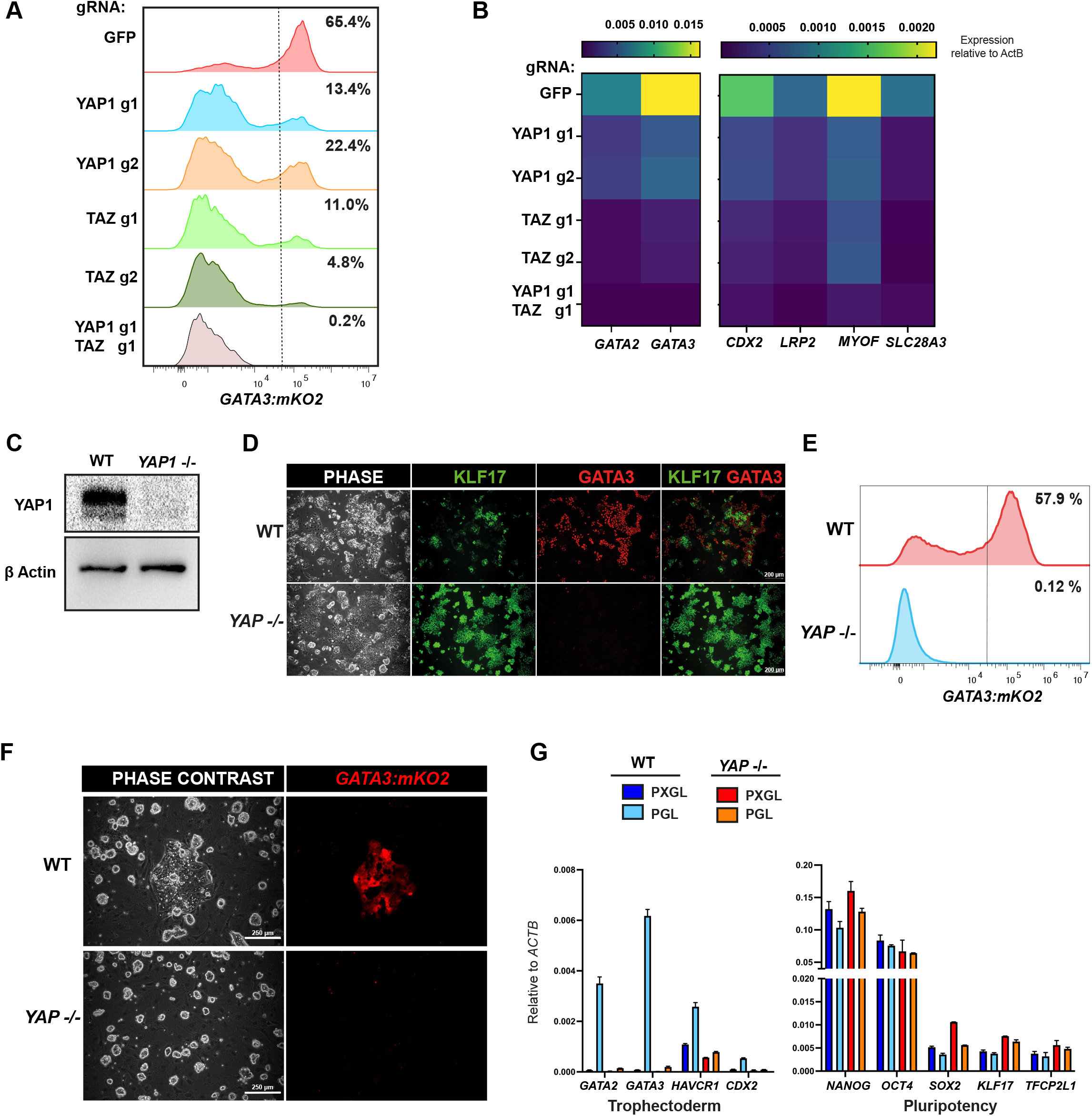
Depletion of YAP1 abolishes TE differentiation and enables sustained self-renewal without XAV. **A**. Flow cytometry analysis for *GATA3:mKO2* reporter in cells transfected with indicated gRNAs. Cells were cultured in PD03 alone for 4 days. **B**. Heatmap of the expression of TE markers in gRNA transfected pools as in 4A. Expression levels are means of duplicate qRT-PCR assays. **C**. Immunoblot of YAP1 in parental and clonal knockout cells. **D**. Immunofluorescence staining of parental and knockout cells for KLF17 and GATA3 following 4 days culture in PL. **E**. Flow cytometry analysis for *GATA3:mKO2* in wild-type and YAP1 knockout cells following 4 days culture in PL. **F**. Spontaneous differentiation to TE-like cells in parental HNES1 *GATA3:mKO2* cultures in PGL. *YAP1*^*-/-*^ cultures remain uniformly dome shaped and undifferentiated. Images were taken 4 days after passage 10 in PGL. **G**. qRT-PCR assay for pluripotency and lineage markers in parental and YAP knockout cultured for long-term in either PXGL or PGL media (10 passages) on MEF layers. RNA was collected at 4 days after passaging. Error bar indicates s.d of PCR duplicates

To examine their phenotype in more detail, we picked and expanded *YAP1* KO cells. Western blotting confirmed absence of YAP1 protein (**Fig. 4C**). When plated in PD03+LIF, *YAP1* KO cells did not up-regulate GATA3 but retained expression of the general pluripotency factor OCT4 and naïve marker KLF17 (**Fig. 4D, 4E**).

We tested whether *YAP1* knockout removes the requirement for XAV in sustaining human naïve PSC cultures. We plated parental HNES1-*GATA3:mKO2* and *YAP1 KO* cells in PGL medium without XAV939 and cultured for over 10 passages. In the parental cultures at each passage areas of *GATA3:mKO2* positive cells appeared after 3-4 days (**Figure 4F**). In contrast, *GATA3:mKO2* positive cells were rarely observed in *YAP1* KO cultures. *YAP1* KO cells maintained naïve markers and showed minimal expression of trophectoderm genes or YAP1/TAZ-TEAD targets. (**Figure 4G**). We conclude that YAP1 mediates human naïve PSC to TE differentiation and that the major effect of XAV939 in human naïve PSC self-renewal is to suppress this action.

These findings clarify the apparent contradiction between the self-renewal requirements of mouse naïve ES cells for GSK3 inhibition, which stabilises β-catenin, and of human nPSCs for tankyrase inhibition, which destabilises *β*-catenin. In mouse, β-catenin prevents the repressor Tcf7l1 from silencing naive pluripotency factors (Hoffman et al., 2013; Martello et al., 2012; Wray et al., 2011). Human nPSCs, however, barely express *TCF7l1* or its major target *ESRRB* (Boroviak et al., 2018; Rostovskaya et al., 2019). Thus, the presence of β-catenin is not essential for human naïve PSC self-renewal. It has alternatively been suggested that degradation of β-catenin via tankyrase inhibition is instrumental for human nPSC propagation (Bayerl et al., 2021). However, our findings demonstrate that deletion of *β*-catenin does not change nPSC dependency on XAV939. The relevant effect of tankyrase inhibition is to stabilise AMOTL2 and reduce YAP nuclear activity. YAP regulation is critical because of the capacity of human nPSCs for TE differentiation, a fate that is not open to mouse ES cells.

YAP has a well-established role in early lineage segregation in the mouse embryo (Nishioka et al., 2009). In outer cells nuclear accumulation of YAP1/TAZ promotes transcription of TEAD target genes that promote TE differentiation, while in inner cells the AMOT complex limits YAP access to the nucleus (Hirate et al., 2013; Leung and Zernicka-Goetz, 2013). nPSCs offer new potential for dissecting mechanisms of early embryogenesis in human. Our study indicates that the key role of YAP1/TAZ in TE differentiation is conserved between species, in line with recent comparative embryology studies (Gerri et al., 2020). Strikingly, although human nPSCs relate to the early epiblast, a developmental stage later than morula where TE lineage specification and segregation initially occurs in embryos, the action of YAP1 in promoting TE differentiation endures. A second component of PXGL (and its forerunner t2iLGö (Takashima et al., 2014)), the aPKC inhibitor Gö6983, further buttresses human naïve PSCs against TE differentiation (Guo et al., 2021). aPKC inhibition restricts establishment of apicobasal polarity, which is also implicated in human embryo TE lineage segregation (Gerri et al., 2020).

Two additional features are worthy of note. Firstly, although AMOT depletion results in TE induction in human nPSCs, it does not do so in the absence of MEK/ERK inhibition. Future investigations will reveal how inhibition of MEK/ERK enables TE lineage specification in human nPSCs while simultaneously suppressing the formative transition. Secondly, we have been unable to expand nPSCs doubly deficient for both *YAP1* and *TAZ*. This suggests that activity of these transcriptional co-factors at low level is necessary for human nPSC propagation and it will be of interest to delineate the specific targets.

## Supporting information

Supplemental Table of Reagents

## ACKNOWLEDGEMENTS

We thank Nia Morris and Francesca Carlisle for providing laboratory assistance. We thank Corin Liddle at the Bioimaging centre, University of Exeter, for imaging support. This research was funded by the Medical Research Council (MRC) of the United Kingdom (MR/P00072X/1). AS is an MRC Professor (G1100526/1).

## AUTHOR CONTRIBUTIONS

Conceptualization, GG, AD; Methodology, AD; Investigation, AD, TH; Writing, AD, GG, AS; Supervision GG, AS

## MATERIALS AND METHODS

### Cell culture

Human nPSCs HNES1-*GATA3:mKO2/*Cas9 (Guo et al., 2021)and niPSCs (Bredenkamp et al., 2019b) were published previously. Cells were maintained without antibiotics and regularly tested negative for mycoplasma by PCR.

Human naive stem cells were propagated in N2B27 with PXGL [1μM PD0325901 (P), 2μM XAV939 (X), 2μM Gö6983 (G) and 10ng/mL human LIF (L)] on irradiated or mitomycin-inactivated MEF feeders as described previously in Bredenkamp et al., 2019b. Cultures were passaged by dissociation with Accutase (Biolegend, 423201) every 3 to 5 days. Rho-associated kinase inhibitor (Y-27632) and Geltrex (0.5μL per cm^2^ surface area; hESC-Qualified, Thermo Fisher Scientific, A1413302,) were added during replating.

### Differentiation assays

Human naive cells were plated in PXGL with Y-27632 on Geltrex. The day after plating, cultures were washed with PBS and transferred to N2B27 with appropriate inhibitors or cytokines for the particular assay. Concentrations of inhibitors/cytokines used were: 1μM PD0325901, 2μM XAV939, 2μM Gö6983 and 10ng/mL human LIF (L). Medium was refreshed every day thereafter.

### Knockout by gRNA plasmid transfection in Cas9 expressing naive cells

gRNA oligos (Supplemental Table 1) were annealed to double-stranded DNA and cloned into a *Piggybac (PB*) vector (CML32) following a U6 promoter. CML32 contains a puromycin resistance gene and a T2A-BFP gene (Guo et al., 2021). gRNA-expression plasmids were transfected together with *PBase* plasmid into HNES1*-GATA3:mKO2* cells using the Neon Transfection system (Invitrogen). HNES1*-GATA3:mKO2* cells are engineered to express *Cas9* constitutively from the *AAVS1* genomic locus (Guo et al., 2021). Following transfection, cells were plated onto feeder-free Geltrex-coated plates in PXGL with Y-27632 for 1 day then exchanged to culture media relevant for the assay. When assessing TE differentiation dependency on PD03 (as in Fig. 3C), cells were transfected and directly plated into PXL or N2B27. Puromycin (0.5 μg/mL) was then applied for at least 3 days to select cells with *PB-gRNA* plasmid integration.

### AMOTL2 overexpression

cDNA encoding the longest isoform of AMOTL2 was amplified by PCR from total cDNA and cloned into a TOPO pENTR/D-TOPO vector using pENTR/D-TOPO cloning kit (Invitrogen). After Sanger sequencing AMOTL2 cDNA was Gateway cloned into a PiggyBac vector behind a CAG promoter. The PiggyBac vector contains a PGK-hygromycin cassette. *PB-CAG-AMOTL2* plasmid was transfected into *GATA3:mKO2/Cas9* reporter cells using the Neon transfection system. Hygromycin selection was applied for 24 hrs to establish stable transgenic cell lines.

### Reverse transcription and real-time PCR

Total cellular RNA was extracted using ReliaPrep kit (Promega, Z6012) and cDNA synthesized with GoScript reverse transcriptase (Promega, A5004) and 3’Race (oligo dT) adaptor primers. TaqMan assays (Thermo Fisher Scientific) and Universal Probe Library (UPL) probes (Roche Molecular Systems) were used to perform gene quantification. A list of UPL primers is provided in Supplemental Table 1.

### Western blotting

Cells were scraped from adherent cultures and lysed with RIPA lysis buffer (150mM NaCl, 50mM Tris pH 8.0, 1% Triton x-100, 0.5% Deoxycholate, 1mM EDTA, 0.1% SDS) supplemented with benzonase, phosphatase and protease inhibitors. Protein concentrations were determined using a 660nm Pierce protein assay. At least 5 μg of protein was mixed with the appropriate amount of sample buffer and DTT (10mM final) and heat-denatured before separation on a 10% SDS-PAGE gel. Following electrophoresis, proteins were transferred onto a PVDF membrane and blocked with 5% bovine serum albumin in Tris-buffered saline containing Tween-20 (0.1%) (TBST) for 1 h at room temperature. Primary antibodies were diluted in blocking solution and incubated with the membrane overnight at 4°C or 2 hrs at room temperature. After washing with TBST, the membrane was incubated with species-specific HRP conjugated antibodies diluted (1:1000) in blocking solution. Novex ECL Chemilumiescent substrate reagent kit was used for developing the membrane and imaging was carried out on the BioRad GelDoc XR+ system.

### Immunofluorescence staining of adherent cells

Adherent cells were washed twice with PBS and fixed with 4% formaldehyde. Cells were permeabilised with 0.3% Triton-X/PBS solution for 15 minutes, and subsequently incubated with 5%BSA/PBS blocking solution for 1 hr at room temperature. Samples were incubated with primary antibodies (Supplemental Table 1) at 1:500-1:1000 dilution in blocking solution either for 1-2 hrs at room temperature or overnight at 4c. After washing 3 time for 15 minutes with 0.1% Triton-X/PBS solution secondary AlexaFluor antibodies were applied at 1:1000 in blocking solution. Cells were washed with 0.1% Triton-X/PBS at least 3 times for 15 minutes. Cells were stained with DAPI. Imaging was performed on a Leica DMI-8 or Zeiss LSM880 in Airyscan mode.

### Flow cytometry analysis

For β-catenin staining, cells were dissociated with TrypLE, and washed twice with PBS, spinning down 300g for 5 minutes. Cells were fixed with 4% formaldehyde solution for 15 mins on a rotator mixer and washed twice with PBS. Cells were blocked in 2% foetal bovine serum (FBS)/PBS for 1 hr at room temperature and incubated with primary antibodies for 1 hr in blocking solution. Cells were washed three times with PBS and incubated with AlexaFluor 488 antibodies in blocking solution for 1 hr at room temperature. Cells were washed 3 times and resuspended with PBS and processed for flow cytometry.

For surface marker SUSD2/CD24 staining, live cells were dissociated with TrypLE, washed, and incubated with directly conjugated antibodies diluted in PBS with 2% FBS for 1 hr at 4°C. Cells were washed and resuspended in PBS.

Flow cytometry was carried out on a CytoFlex cytometer (Beckman Coulter) with analysis using FlowJo software. Gating was kept consistent across all samples in each experiment. TOPRO3 staining was used to exclude the dead cell population.

