## Supplemental figures and legends for "Suppression of YAP Safeguards Human Naïve Pluripotency"

### SUPPLEMENTAL FIGURE LEGENDS

**Supplemental Figure 1: Deletion of β-catenin in naïve PSC self-renewal and TE differentiation**

**S1A**. Schematic of strategy for generating and validating *CTNNB1*$^{-/-}$ human nPSCs

**S1B**. Immunofluorescence staining for β-catenin in wild-type and clonal *CTNNB1*$^{-/-}$ nPSCs. Scale bar: 20 µm.

**S1C**. Phase image showing morphology of wild-type and *CTNNB1*$^{-/-}$ nPSCs following 3 and 4 days of culture in PXGL on MEF. *CTNNB1*$^{-/-}$ display reduced cell-cell adhesion at day 4. Scale bar: 50 µm.

**S1D.** qRT-PCR for pluripotency markers in wild-type and *CTNNB1*$^{-/-}$ cells.

**S1E**. Immunofluorescence staining for naïve marker KLF17 in wild-type and CTNNB1$^{-/-}$ nPSCs cultured in PXGL on MEF. Scale bar: 100 µm.

**S1F**. Immunofluorescence staining for β-catenin and GATA3 in GFP gRNA and CTNNB1 gRNA transfected pools following withdrawal of XAV for 3 days from PXGL. Scale bar: 100 µm.

**S1G.** Immunofluorescence staining for Plakoglobin and GATA3 in wild-type and *CTNNB1*$^{-/-}$ cells following culture in PL for 3 days. Scale bar: 100 µm.

**Supplemental Figure 2: Expression of AMOT family and YAP-TEAD targets**

**S2A.** FPKM expression values of AMOT family genes derived from scRNA-seq datasets for early human embryo. Boxplots were generated using GRAPPA (Boroviak et al., 2018). The midline identifies the median, whiskers the first and third quartile expression levels, and each dot is an individual cell.

**S2B**. Complete immunoblot for Fig. 2C showing AMOTL2 expression in indicated conditions 24 hrs after transfer from PXGL.

**S2B**. Immunofluorescence staining for AMOTL2 in nPSCs 48 hrs after XAV withdrawal from PXL cultures. Scale bar: 100 µm.

**S2D**. Western blot showing AMOTL2 expression in indicated conditions 48 hrs after transfer from PXGL.

**S2E**. Expression of YAP/TEAD target genes in niPSCs cultured in PXGL or PGL for 3 days.

**Supplemental Figure 3: Genetic manipulation of AMOTs and TE differentiation**

**S3A.** Flow cytometry analysis for *GATA3:mKO2* reporter in cells transfected with indicated AMOT gRNAs. Cells were kept in PXGL.

**S3B.** Western blot of AMOTL2 6 days following transfection with GFP or AMOTL2 gRNA. Cells were cultured in PXGL before protein collection.

**S3C**. Immunofluorescence staining for YAP1 and GATA3 after transfection with GFP or AMOT/AMOTL2 gRNAs and culture for 5 days in PXL. Scale bar: 200 µm.

**S3D.** qRT-PCR for trophectoderm markers for parental and AMOTL2 OE cells in PL 3 days after transfer from PXGL. Error bars indicate s.d. of PCR duplicates.

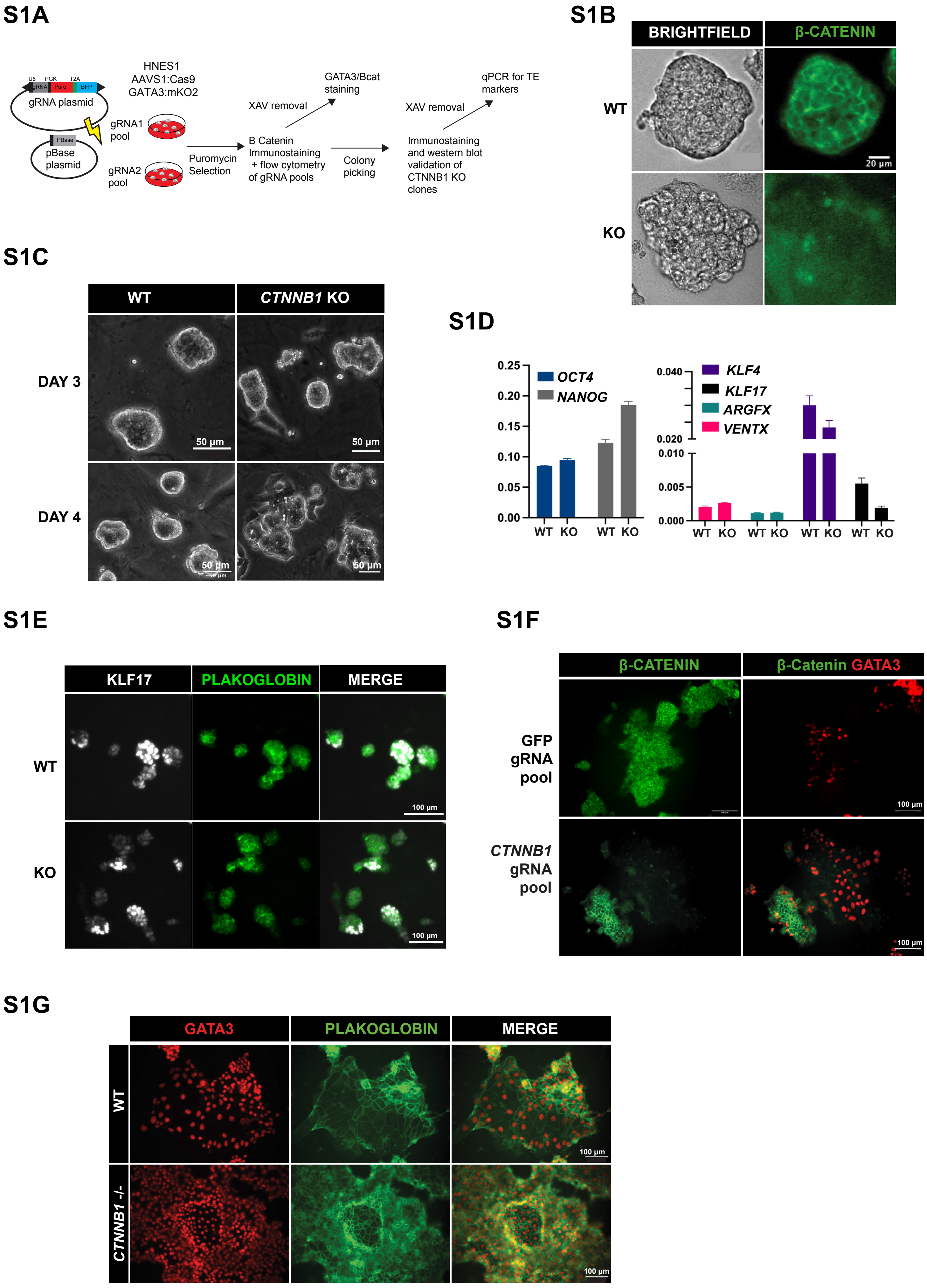


**Supplement Figure 1**

S2A

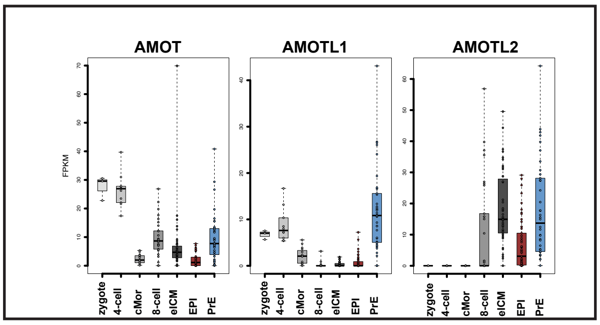


S2B

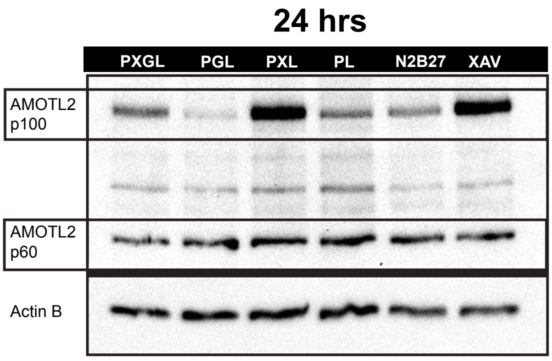


S2C

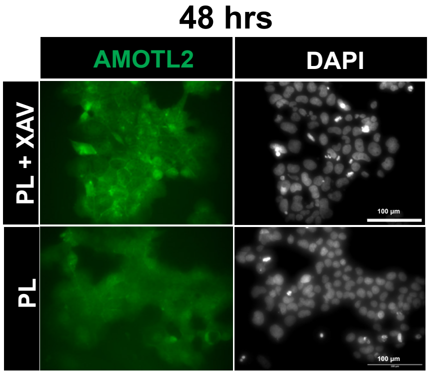


S2D

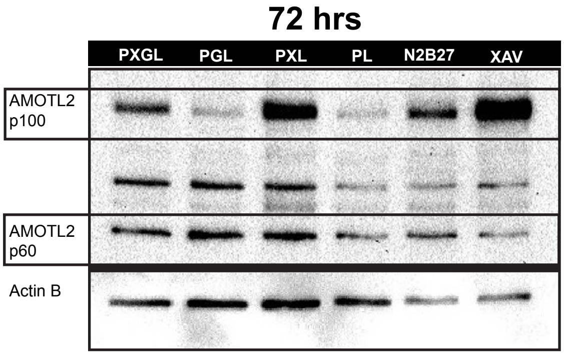


S2E

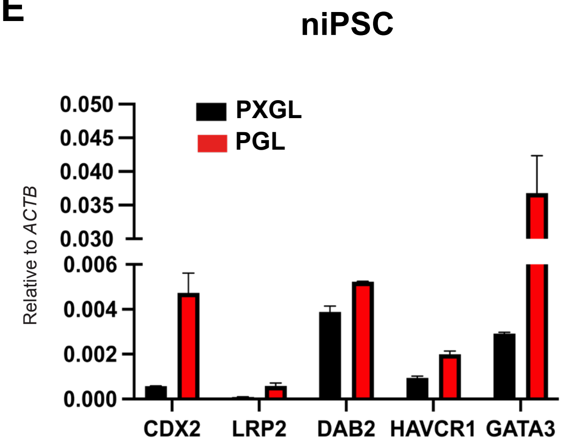


### Supplement Figure 2

S3A

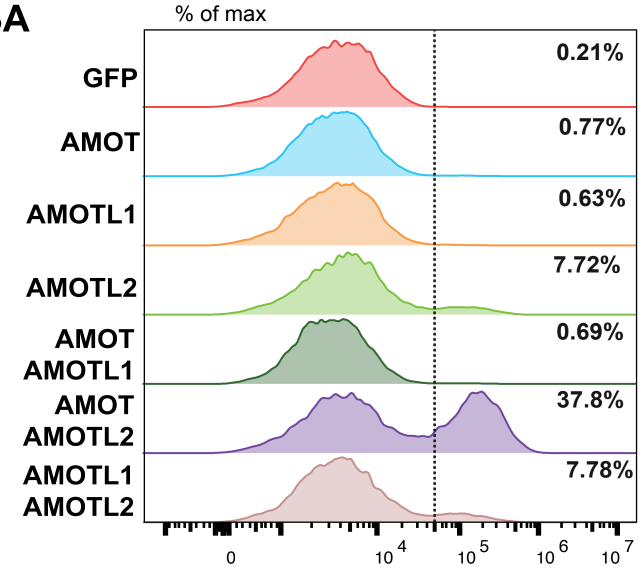


S3B

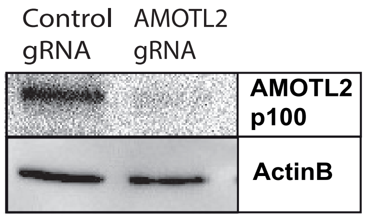


S3C

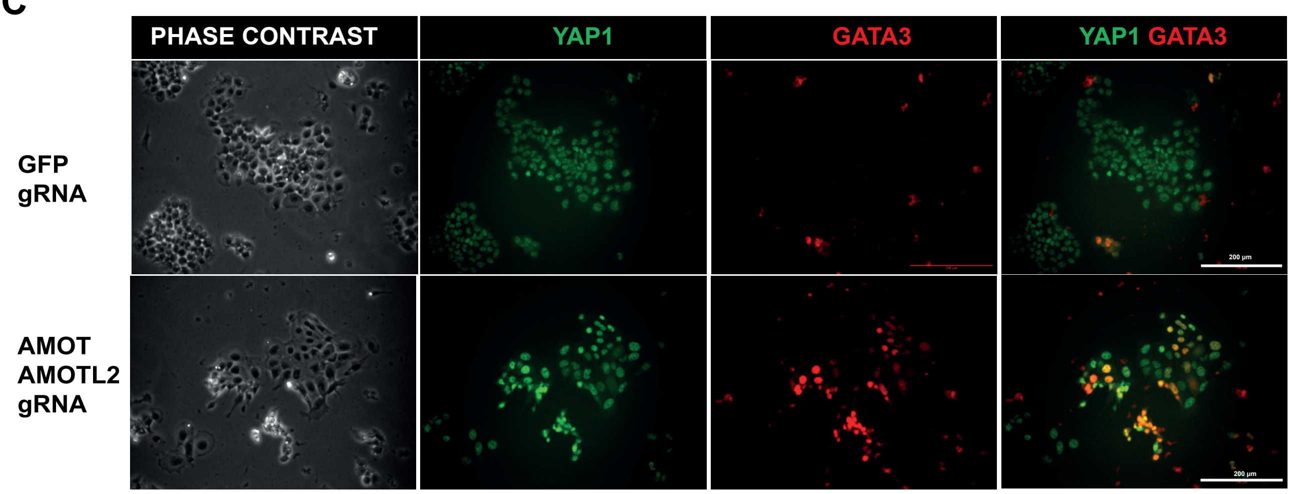


S3D

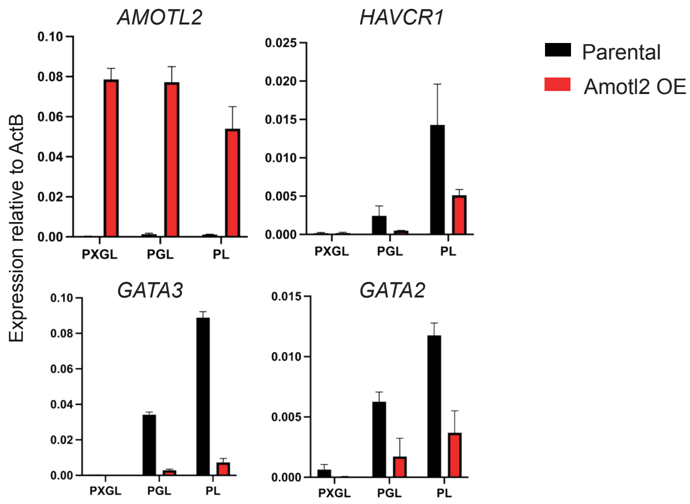


### Supplement Figure 3
